# Closed-loop sonothermogenetic control of CAR T cells for metronomic brain cancer therapy

**DOI:** 10.1101/2025.04.23.650339

**Authors:** Ali Zamat, Chulyong Kim, Shwetha Sridhar, Sydney Fabrega, Riya Sen, Noah Campbell, S. Abbey Oliver, Zizhen Zha, Chloé Thiveaud, Elif Kulaksizoglu, Meredith Brienen, Hideho Okada, Graeme F. Woodworth, Costas Arvanitis, Gabriel A. Kwong

## Abstract

Achieving durable CAR T cell responses against primary brain tumors and metastases requires strategies that enable intracranial control of therapy to overcome the barriers of solid tumor treatment without compromising safety. Here, we show that closed-loop sonothermogenetics enables remote regulation of CAR T cell therapeutic activity through the intact skull. Using MR-guided focused ultrasound with closed-loop temperature feedback, we modulate CAR T cells engineered with a genetically encoded thermal bioswitch to achieve metronomic activation in the brain without lasting adverse effects on healthy brain tissue. In murine models of brain cancer, metronomic production of NKG2D T cell engagers by intratumoral CAR T cells overcomes antigen heterogeneity in breast cancer brain metastasis and myeloid-derived immunosuppression in glioblastoma to drive antitumor responses. Our findings support the use of closed-loop sonothermogenetics for spatial and temporal control of CAR T cell therapies targeting solid brain tumors.

## Introduction

CAR T cell therapy for brain and central nervous system tumors is rapidly advancing, with multiple clinical trials demonstrating acceptable safety profiles and early positive responses in secondary endpoints for neuroblastoma(*1*), glioblastoma(*2–5*), and diffuse midline glioma(*6, 7*). However, achieving durable clinical benefit will require strategies to overcome the challenges posed by solid tumors, such as immunosuppression and cancer cell antigen heterogeneity, without elevating safety risks (*8–10*). Approaches aimed at enhancing spatial or temporal control of CAR T cell activity have shown promise to improve treatment of brain tumors. For instance, localized delivery methods – such as intracerebroventricular or intracavitary infusion of CAR T cells – have demonstrated increased potency and tolerability compared to intravenous administration (*6*), using local and repeated administration of low CAR T cell doses to regulate therapy (*4, 7, 11*). These studies underscore the potential of CAR T therapy to target intracranial tumors and the importance of locally regulating CAR T cell function to minimize dose-related toxicities without compromising therapeutic potency (*10*).

To regulate CAR T cell activity, a range of strategies have been developed to provide spatial and/or temporal precision (*12*). Among them, small molecule–regulated systems, including ON- and OFF-switch designs, offer reversible and dose-tunable control of CAR T cell function (*13–16*). Likewise, adapter-based platforms, including split, universal, and programmable (SUPRA) CARs, and other switchable receptors, enable modular targeting and titratable functional activity (*17–19*). Although these strategies provide temporal control, their dependence on systemically administered molecules limits spatial specificity and may pose challenges for brain-targeted applications due to the blood-brain barrier(*20*). Logic-gated architectures, such as synNotch-based CAR T cells, can address these challenges by enabling localized expression of CARs following the detection of brain-specific antigens; however, they offer limited temporal regulation once activated (*21–23*). To achieve both spatial and temporal control, CAR T cells designed with genetically encoded thermal bioswitches have emerged as a promising approach, allowing thermal targeting of tissues to a well-tolerated temperature of 42°C to control the magnitude, duration, and location of transgene expression *in vivo (24–27)*. However, despite the potential of thermogenetic control to potentiate brain tumor therapy, achieving safe, repeatable, and effective transcranial control of CAR T cell function remains a major challenge.

Magnetic Resonance Imaging-guided Focused Ultrasound (MRgFUS) presents distinct advantages to enable sonothermogenetic control of CAR T cell activity in the brain. Notably, it can noninvasively target multiple small tissue volumes deep within the brain through an intact skull using either multi-element phased arrays(*28*) or holographic lenses(*29*). Moreover, it can be integrated with MR temperature imaging (MRTI) to provide real-time monitoring and control of temperature elevation in the focal region (*30*). In preclinical studies using extracranial tumors as a model system, MRgFUS has shown promise in regulating CAR T cell therapeutic function(*24*). Clinically, MRgFUS is approved and used for minimally invasive thermoablative procedures to treat neurological diseases (*31–33*). While all these characteristics offer a clear path to translation, achieving the precise temperature control required for repeated activation of CAR T cells without inducing adverse effects on healthy tissue is not trivial, particularly at the skull-brain interface, where the energy absorption of the skull is significantly higher than brain tissue(*28*). Previous studies targeting the brain either did not incorporate MR thermometry(*27*), or used a closed-loop control system that could not maintain temperature within a narrow tolerance (< 0.5°C) (*30*). The absence of precise thermal control poses a significant barrier to clinical translation, as temperatures below 42°C may result in subtherapeutic effects while exceeding this threshold can cause damage to brain tissue.

In this study, we engineered CAR T cells with a thermal bioswitch and employed transcranial MRgFUS to achieve closed-loop sonothermogenetic activation, enabling remote spatiotemporal control of CAR T cell function within the brain (**Fig. 1a-b**). We show that close-loop control of ultrasound pulses can locally and rapidly elevate and maintain brain tumor temperatures at 41.5 ± 0.1°C for thermal bioswitch activation without causing lasting adverse effects on either the targeted tissue or the CAR T cells. We show that metronomic activation of CAR T cells engineered to produce NKG2D T cell engagers potentiates αHER2 CAR T cell therapy against breast cancer brain metastasis by overcoming antigenic heterogeneity, or αEGFRvIII CAR T cell therapy against GBM to mitigate myeloid derived suppressor cell (MDSC) mediated immunosuppression. Collectively, our findings demonstrate that sonothermogenetic control of CAR T cell function can boost antitumor potency, overcome resistance mechanisms, and is well-tolerated for the treatment of intracranial malignances.

**Fig. 1.**
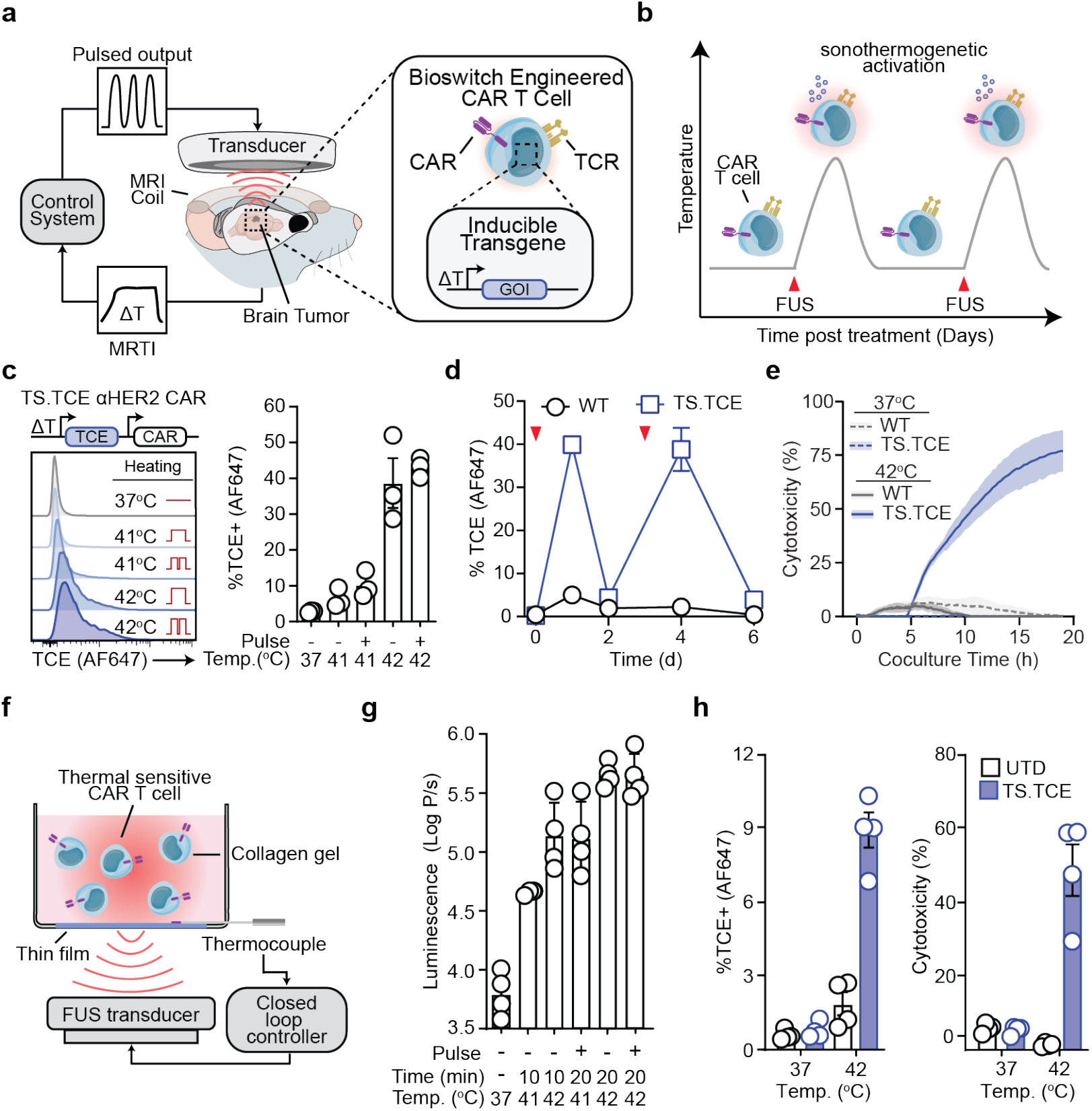
Closed-loop FUS enables sonothermogenetic control over CAR T cell activity. (**a**) Schematic representation of sonothermogenetic control of thermal sensitive T cells by MRgFUS across an intact skull. (**b**) Representative schematic demonstrating the transient and repeatable nature of thermal switch activity to enable metronomic dosing. (**c**) Representative flow staining and quantitation of TCE expression following 20-minute heating at either 41°C or 42°C with continuous or pulsed heating strategy and (**d**) quantification of transient TCE production of following two thermal treatments. (**e**) Quantification of cytotoxicity against HER2-MDA-MB-468 following coculture of WT or engineered T cells following thermal treatment at 37°C or 42°C. (**f**) Schematic representing *in vitro* closed-loop FUS setup to evaluate the effects of FUS-mediated hyperthermia on transgene production of CAR T cells. (**g**) Luminescence of TS.Fluc aHER2 CAR T cells following FUS-mediated hyperthermia at indicated parameters quantified by IVIS. (**h**) TCE expression (left) and cytotoxicity (right) against HER2-MDA-MB-468 tumor cells following FUS mediated hyperthermia for 10 minutes at indicated temperatures. Mean ± s.e.m. depicted, n= 3-4 independent wells.

## Results

### Metronomic control of CAR T cell activity via closed-loop sonothermogenetics

Primary human αHER2 CAR T cells were genetically modified with a previously described thermal bioswitch (TS) (*25*) to control the production of an NKG2D T cell engager (TCE). NKG2D TCEs allow CAR T cells to target stress-induced antigens present on cancer cells in addition to the CAR-targeted antigen, thereby expanding their reactivity against tumor cells (*25*). Heating TS.TCE CAR T cells with a thermal cycler at 41°C or 42°C induced a temperature-dependent increase in TCE surface staining under both continuous and pulsed (66% duty cycle) conditions (**Fig. 1c**). TCE expression returned to baseline levels of wildtype (WT) cells within two days and remained inducible upon repeated heating (**Fig. 1d**), demonstrating the potential of metronomic dosing to mitigate exhaustion and sustain CAR T cell function (*34–36*). When heated and cocultured with HER2-tumor cells, only TS.TCE αHER2 CAR T cells exhibited cytotoxicity, confirming that thermal activation is required to redirect CAR function (**Fig 1e**).

We confirmed the use of FUS for closed-loop sonothermogenetic activation of CAR T cells within a collagen tissue phantom to minimize non-thermal acoustic effects (**Fig. 1f**). We delivered FUS exposures using continuous wave sonication at a referenced peak negative pressure of ~0.8 MPa, corresponding to an estimated spatial-peak intensity of ~21.3 W/cm^2^ — sufficient to achieve ~41 – 42°C in the collagen gel. Under closed-loop feedback control, FUS activation of TS.Fluc αHER2 CAR T cells at 42°C induced a ~79-fold increase in reporter transgene expression (**Fig. 1g, S1**). FUS-mediated heating at 41°C or 42°C for 20 minutes had no significant effect on CAR T cell cytotoxicity against HER2+ MDA-MB-468 (**Fig. S2a-b)**. However, IFNγ production was mildly reduced at 42°C while secretion at lower temperatures remained comparable to levels observed at 37°C (**Fig. S2c**). Closed-loop sonothermal activation of TS.TCE CAR T cells at 42°C triggered TCE expression which exhibited significant cytotoxicity against HER2-tumor cells, while unheated or WT cells remained inactive (**Fig. 1h**). Taken together, these results demonstrate that closed-loop FUS-mediated heating can activate thermal bioswitches, enabling the titratable expression of TCEs and the regulation of cytotoxic function by engineered CAR T cells.

### Closed-Loop MRgFUS enables spatial and temporal sonothermal control for intracranial hyperthermia

Due to the stringent temperature control requirements for robust and repeated bioswitch activation and not overriding the thermal regulation of the brain — a highly sensitive organ requiring control over temperature distribution, spatial targeting, and temporal regulation — the FUS system needs careful tuning. In our previous investigations, we optimized the FUS transducer design for transcranial hyperthermia (*30*). Here, we further developed a closed-loop MRgFUS system integrating a Proportional-Integral-Derivative (PID) controller to enable tight control of the focal temperature for pulsed activation of the bioswitch in the brain through intact skull. This system is evaluated for its ability to achieve localized hyperthermia in a breast cancer brain metastasis (BCBM) mouse model (**Fig. 2a-b**). The setup combines real-time MRI thermometry imaging (MRTI) for temperature feedback with a closed-loop control algorithm to adjust FUS input dynamically. Tumor regions were identified using T2-weighted MRI, and precise thermal targeting was achieved using feedback correction from MRTI data. To assess the robustness of our system, we compared two control strategies — bistate (**Fig. 2c**) and PID (**Fig. 2g**) — to maintain the target temperature of 41.5 °C. The bistate controller used binary input adjustments to control voltage output (**Fig. 2d**) to approximate the desired temperature output (**Fig. 2e-f**), while the PID controller continuously fine-tuned the input voltage (**Fig. 2h**) based on error minimization to tightly control temperature output and distribution (**Fig. 2i-j**). An analysis of temperature profiles over a 20-minute FUS hyperthermia session (66% duty cycles) at 41.5 °C (**Fig. 2e, 2i, Fig. S3**) demonstrated that while both controllers reached similar average target temperatures (bistate: 41.4°C vs. PID: 41.5°C), the PID controller exhibited substantially lower deviation values (ΔT < 0.1°C vs. 0.5°C). These findings demonstrate that the PID-based closed-loop system effectively induces localized hyperthermia with temperature control, maintaining stability within a narrow range. This enhancement improves the MRgFUS system’s capability for developing robust transcranial hyperthermia protocols and underscores its potential for thermal applications *in vivo*.

**Fig. 2.**
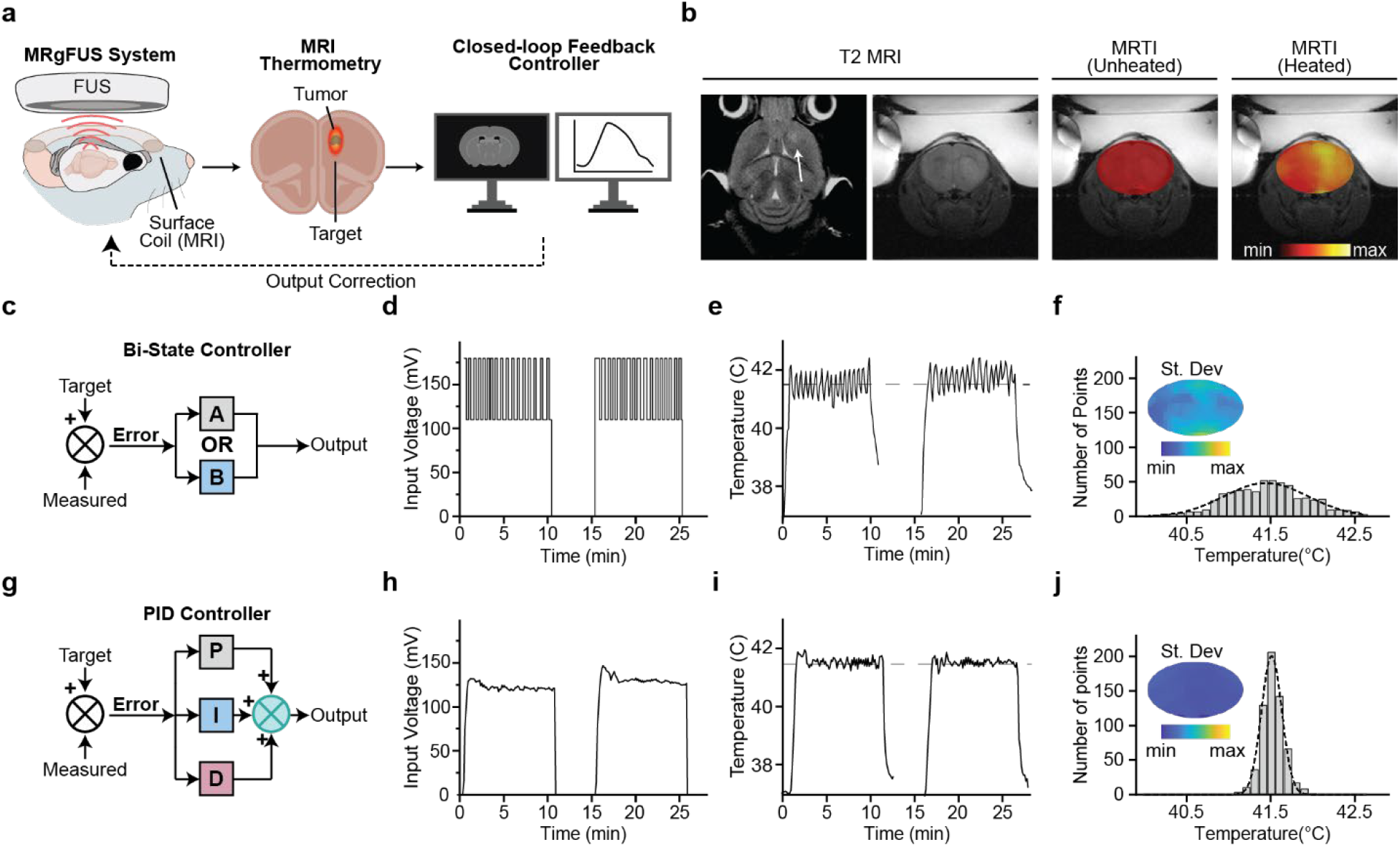
PID-Based Closed-Loop MRgFUS enables precise thermal control for transcranial hyperthermia. (**a**) Schematic of MRgFUS system with closed loop control system. (**b**) Representative images of T2-weighted MRI and fused MRTI. (**c**) Schematic representation of Bi-state controller. (**d**) Representative power output from Bi-state controller. (**e**) Temperature profiles from bi-state controller and (**f**) histogram of temperature variance bi-state controllers. (**g**) Schematic representation of PID controller. (**h**) Representative power output from PID controller. (**i**) Temperature profiles from PID controller and (**j**) histogram of temperature variance bi-state controllers.

### Healthy brain parenchyma tolerates mild hyperthermia

We assessed the tolerability and tissue response to closed-loop controlled localized hyperthermia in the brain. To establish the optimal heating conditions for an *in vivo* mouse model, we focused on a 10-minute hyperthermia treatment at 41.5 °C (67% duty cycle), identified as providing maximal thermal tolerance. Brain sections were analyzed at 2, 7, and 14 days post-treatment (**Fig. 3a**), focusing on markers of astrocyte activation (GFAP), microglial activation (Iba-1), and neuronal degeneration (Fluoro-Jade). MRI revealed mild edema at day 2, which resolved by day 5 (**Fig. 3b**) We observed a significant increase in GFAP and Iba-1 staining at 2 days post-treatment, indicating transient astrocyte and microglial activation, which persisted over several days (**Fig. 3c**). Notably, Fluoro-Jade staining showed no evidence of neuronal degeneration, and Hematoxylin and Eosin (H&E) staining confirmed the absence of tissue damage in the treated regions.

**Fig. 3.**
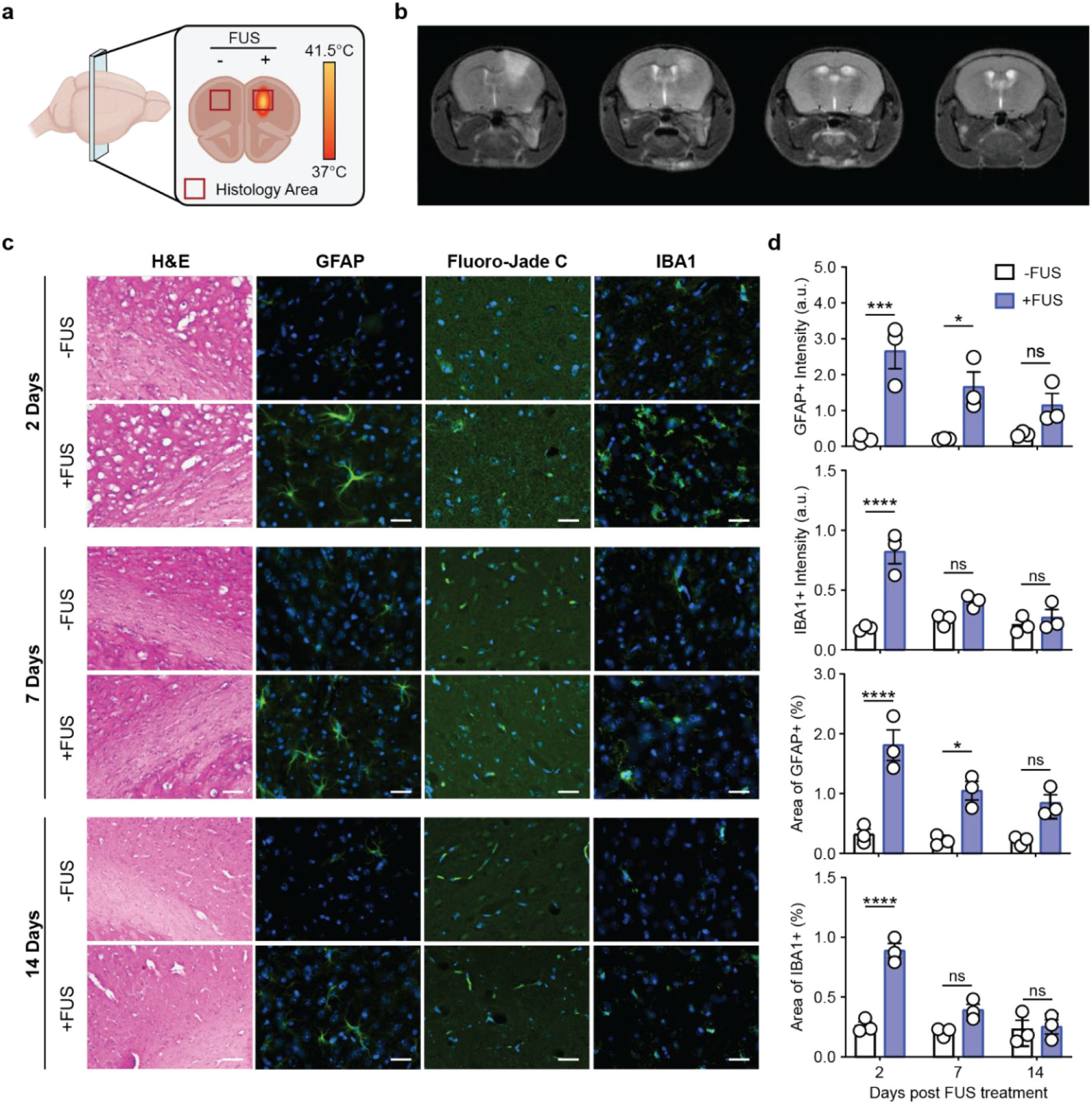
Brain tissue response to transcranial hyperthermia is reversible and well tolerated. (**a**) Schematic showing the brain region safety was used for assessing thermal damage. (**b**) T2-weighted MRI showing edema that dissolves after one week. (**c**) and (**d**) Quantification of microglia and astrocyte activation, demonstrating that they return to baseline 1 - 2 weeks after sonication. (**e**) Representative images of GFAP, IBA-1, Fluoro-Jade and H&E staining at different time points after sonications. For all data n=3.

Quantitative analysis of fluorescence intensity further validated that the levels of GFAP and Iba-1 in the treated regions were significantly elevated compared to the untreated contralateral regions at day 2 (**Fig. 3d**). However, these levels declined over time, returning to non-significant differences compared to baseline. By day 7, Iba-1 expression was indistinguishable from untreated regions, and by day 14, GFAP levels showed no significant difference from baseline. This temporal analysis demonstrates the MRgFUS system safely induces localized hyperthermia without causing long-term neuroinflammation or neuronal damage. Taken together, these experimental findings confirm that the PID-based closed-loop MRgFUS system, operating at an optimal frequency of 1.7 MHz, can safely and effectively induce localized thermal stress in brains of mice without triggering sustained neuroinflammatory or neurodegenerative responses.

### Transcranial activation of CAR T cells is transient, repeatable, and localized

We next applied MRgFUS in murine brain tumor models to noninvasively activate intracranial CAR T cells. We delivered primary human TS.Fluc αHER2 CARs intratumorally via stereotactic injection and targeted the site with sonothermal heating, maintaining tumor temperatures at 41.5 °C (**Fig. 4a**). Luminescence imaging revealed a ~10 fold increase in reporter expression within tumors compared to unheated controls (**Fig. 4b**). Given the transient nature of the thermal bioswitch, and the safety of the MRgFUS-mediated heating, we applied a second round of sonothermal activation, inducing another peak in transgene expression, demonstrating the feasibility of *in vivo* metronomic dosing (**Fig. 4c**). Spatial and temporal control over CAR T cell activity may prevent systemic activation and limit on-target, off-tumor toxicity driven by antigen expression on healthy tissues (*37, 38*). To confirm that sonothermal activation was spatially confined, we analyzed luminescent signal across multiple organs. Bioswitch activity was only detected within the tumor site in heated cohorts, while the liver and spleen showed no significant activation (**Fig. 4d-e**). Taken together, MRgFUS enables localized CAR T cell activation deep within the brain with minimal off-target gene expression to establish sonothermogenetic control over CAR T cells as a noninvasive method for transient, repeatable, and spatially confined immunotherapy.

**Fig. 4.**
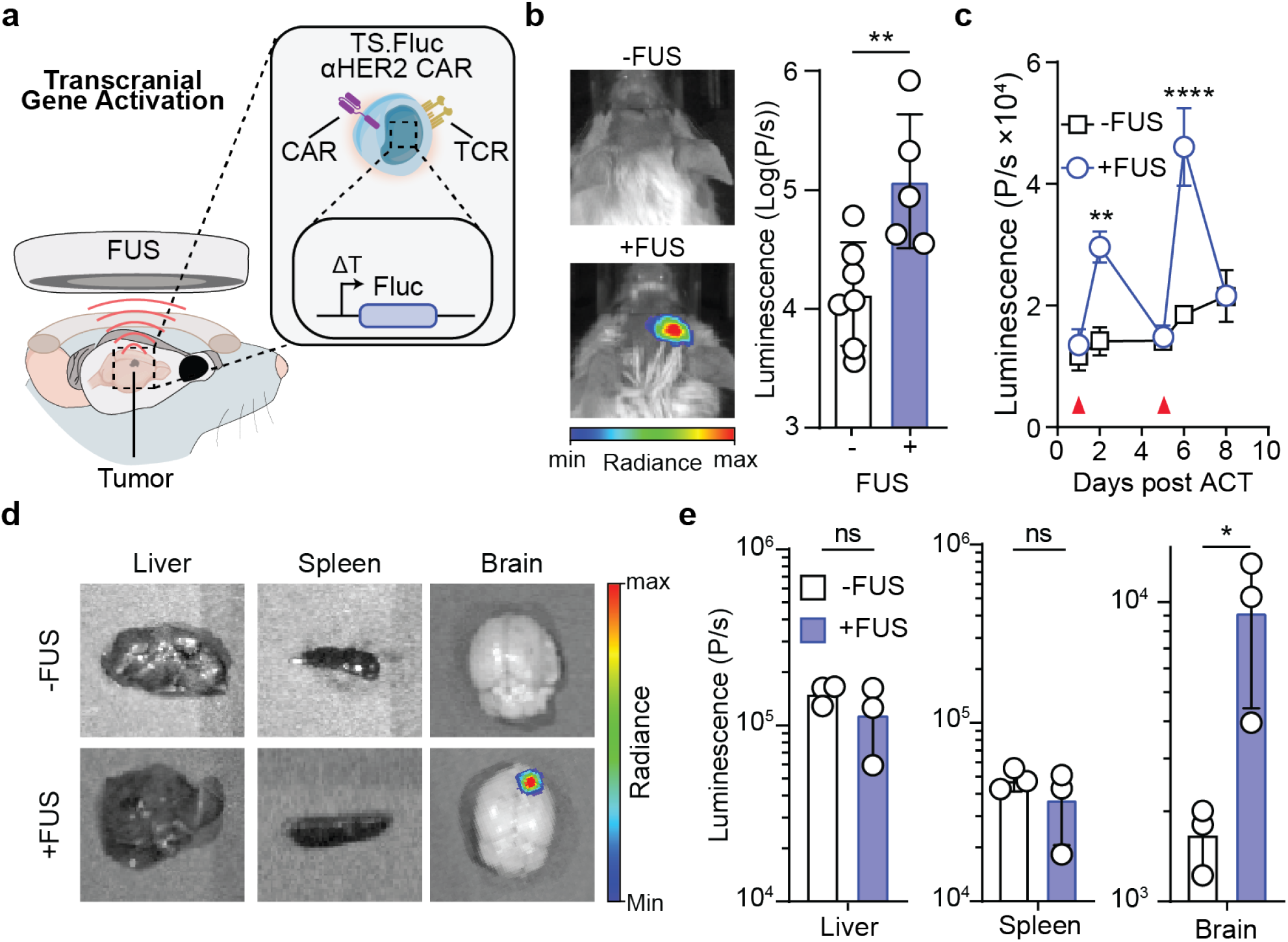
Transcranial activation of CAR T cells is transient, repeatable, and localized. (**a**) Schematic representation of MRgFUS setup with brain-tumor bearing mice and representative in vivo heating strategy. (**b**) Representative image and quantification of luciferase signal from intratumoral TS.Fluc aHER2 CAR T cells 10 hours post MRgFUS-mediated heating. Student’s two-tailed T test, **p < 0.01, mean ± s.e.m. depicted, n=5-7 mice. (**c**) Transient and repeatable activation of thermal sensitive CAR T cells following MRgFUS mediated heating for 10 minutes at 41.5°C (red triangle) as measured by IVIS expression of luciferase. Mixed-effects analysis, **p < 0.01, ****p < 0.0001, mean ± s.e.m. depicted, n=3-5 mice. (**d**) Representative images and (**e**) quantification of liver, spleen, and brain 10 hours post heating, organs were isolated and luciferase signal quantified. Student’s two-tailed T test, *p < 0.05, **p < 0.01, ****p < 0.0001, ns = nonsignificant, mean ± s.e.m. depicted, n=3 mice.

### Sonothermogenetic production of TCEs mitigates antigen escape in heterogenous BCBM tumors

Antigen heterogeneity of brain tumors limits the efficacy of immunotherapies that target a single tumor-specific antigen (*5, 39*) resulting in transient responses and tumor escape (*40–42*). We postulated that sonothermogenetic control could be used to noninvasively drive the intratumoral production of NKG2D TCEs to redirect CAR T cell cytotoxicity towards CAR antigen-negative tumor cells and prevent antigen escape. To test this, we inoculated mice with 75% HER2+/Fluc- and 25% HER2-/Fluc+ MDA-MB-468 cells to model heterogenous breast cancer brain metastasis (BCBM) (**Fig. 5a, Fig. S4a**) and confirmed that sonothermal treatment alone did not alter tumor composition (**Fig. S4b**). To assess therapeutic efficacy, we monitored tumor burden in BCBM mice that received TS.TCE αHER2 CARs (TS.TCE +FUS), TS.Rluc αHER2 CARs as a control for TCE activity CARs (TS.Rluc +FUS), and mice that did not receive therapy (WT-FUS). While sonothermal activation of TS.Rluc αHER2 CARs significantly slowed tumor growth relative to untreated animals (**Fig. b-c**), as measured by MRI quantification of tumor sizes, residual HER2-tumor cells were not eliminated and rapidly expanded, as indicated by bioluminescent imaging (**Fig. 5d**). By contrast, no tumor burden was detected either by MRI or BLI in the cohort that received TS.TCE αHER2 CARs (**Fig. S5**) (**Fig. 5b-e**), achieving significantly extended survival compared to controls (**Fig. S6**). Six of eight mice survived beyond 100 days post-transfer, whereas all control mice succumbed by day 30 (WT-FUS vs. TS.TCE +FUS: p = 0.0003; TS.Rluc +FUS vs. TS.TCE +FUS: p = 0.0191). Taken together, these findings demonstrate that sonothermal activation of CAR T cells enables localized TCE production to redirect CAR T cell cytotoxicity to overcome antigen heterogeneity and achieve durable survival in a heterogeneous BCBM model.

**Fig. 5.**
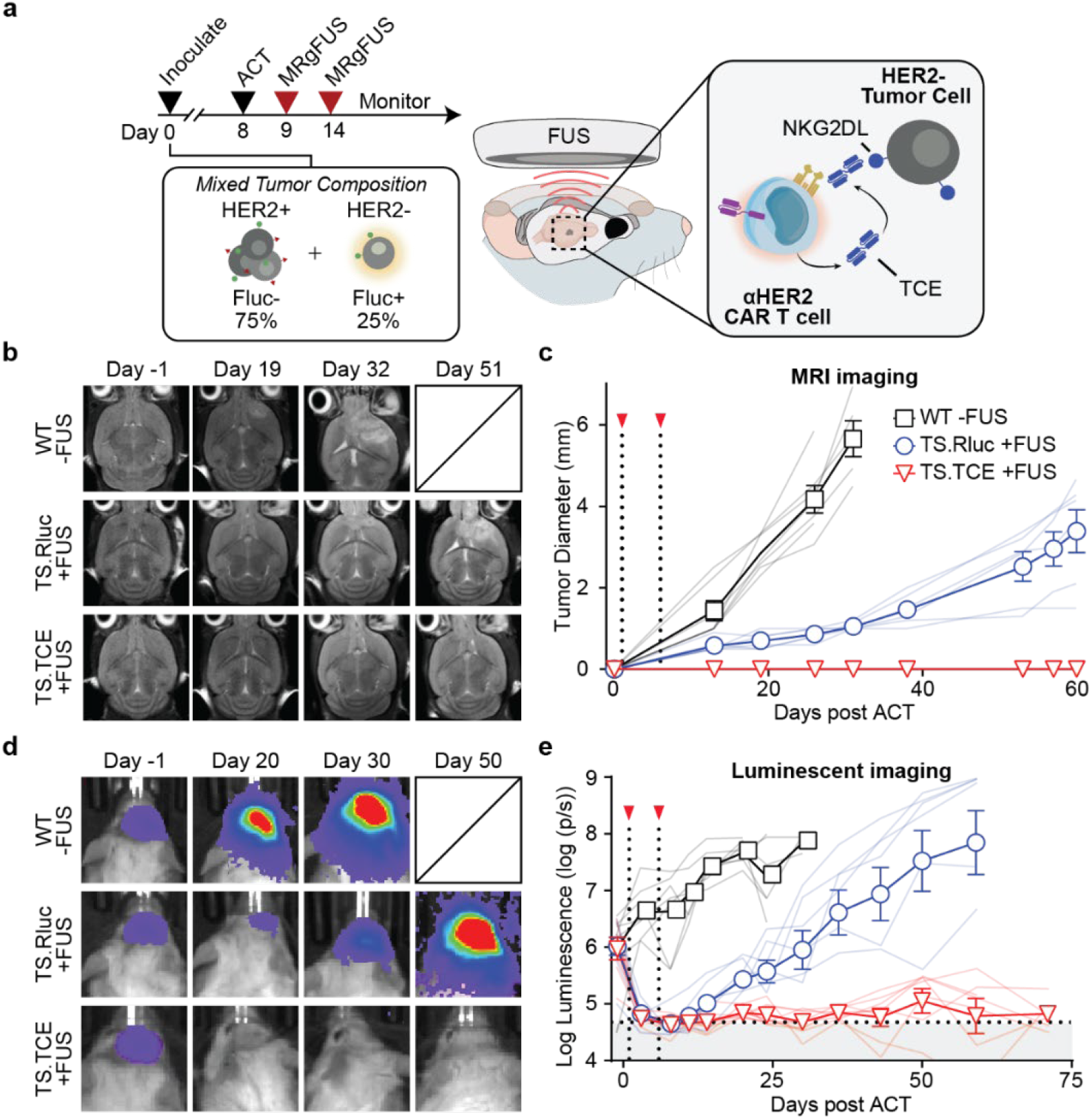
Metronomic production of TCEs mitigates antigen escape in heterogenous BCBM tumors. (**a**) NSG mice were inoculated with heterogenous tumors comprised of a 75% HER2+ tumor mixture of MDA-MB-468 tumor cells. Schematic representation of TCE production following two MRgFUS-mediated thermal treatments to redirect TS.TCE aHER2 CAR T cells towards HER2-tumors. (**b**) Representative MR images and (**c**) quantification following treatment. (**d**) Representative IVIS images and (**e**) quantification of HER2-Tumor burden following treatment, mean ± s.e.m. depicted, n=6-8 mice.

### Local mTCE production remodels the immunosuppressive TME in favor of CAR T cell therapy

Despite T cell infiltration, elevated levels of MDSCs in glioblastoma contribute to immune suppression and correlate with poor prognosis (*43*), making their depletion a promising therapeutic strategy (*44–47*). We therefore tested whether sonothermal production of a murinized NKG2D T cell engager (mTCE) by αEGFRvIII CAR T cells could resist MDSC-mediated suppression and enhance antitumor response (**Fig. 6a**). To first confirm that NKG2D mTCEs can mitigate MDSC suppression, conditioned media from producer cells (mBTE.293T) was added to a co-culture of WT T cells with bone marrow derived MDSCs (**Fig. S7**). At all MDSC to WT T cell ratios tested, we found that mTCE-conditioned media increased T cell increased proliferation, as measured by cell trace violet (CTV) CTV dilution, and activation, as indicated by CD25 expression (**Fig. 6b-c**), demonstrating that mTCEs can overcome MDSCs-mediated immunosuppression.

**Fig. 6.**
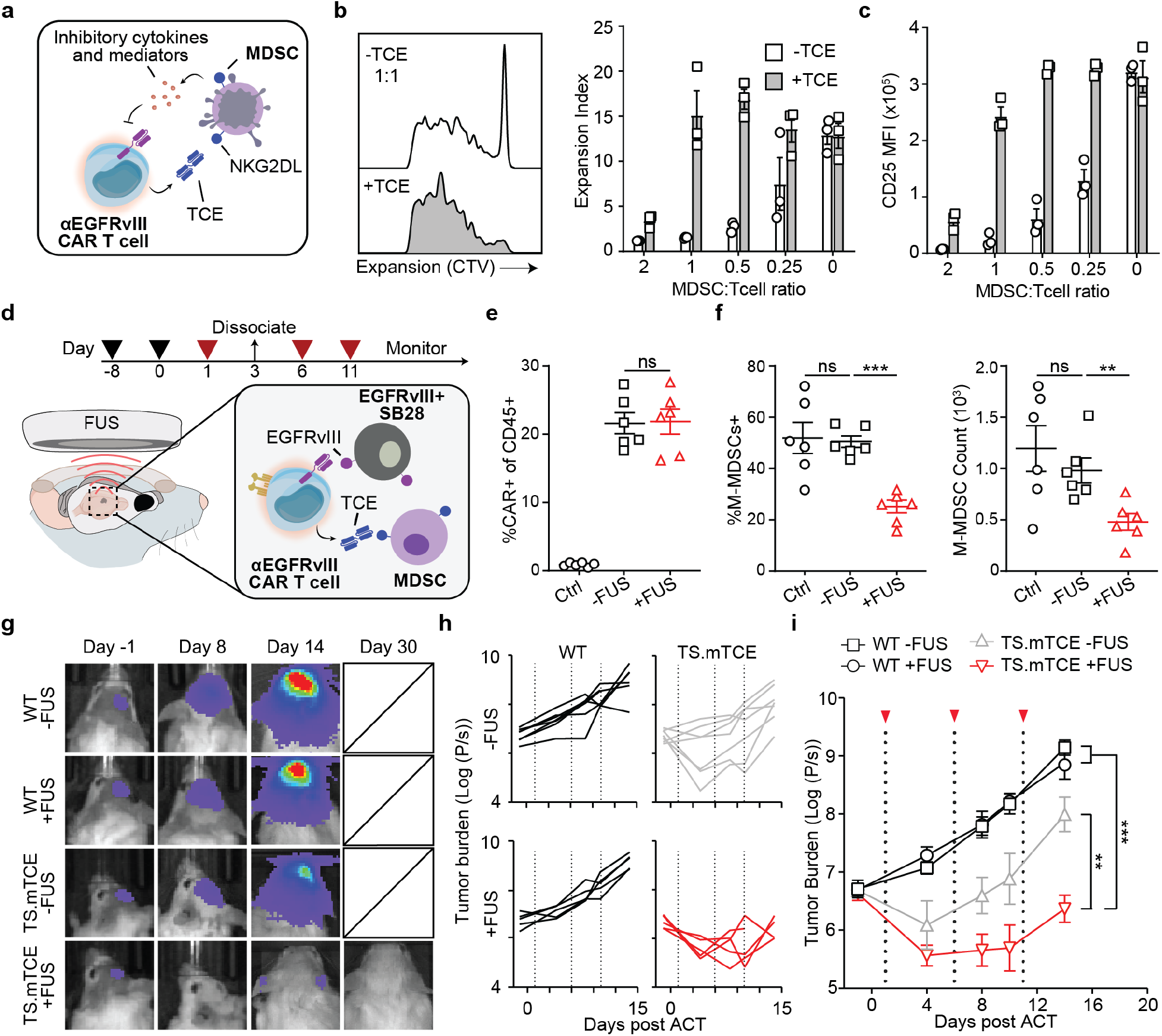
Local mTCE production remodels the immunosuppressive TME in favor of CAR T cell therapy. (**a**) Schematic representation of TCEs redirecting CAR T cell activity towards MDSCs. (**b**) Representative histogram and quantification of expansion index by CTV staining of T cells following coincubation with MDSCs at various ratios. (**c**) Quantification of T cell activation by CD25 staining at varied MDSC:T cell ratios in mTCE conditioned media. (**d**) Schematic of *in vivo* depletion of MDSCs by TS.mTCE EGFRvIII CAR T cells following MRgFUS mediated heating. Quantification of (**e**) percent CAR+ cells and (**f**) percent and count of M-MDSCs cells following MRgFUS treatment of tumor bearing mice. One-way ANOVA with multiple comparisons, **p < 0.01, ***p < 0.001, ns = nonsignificant, mean ± s.e.m. depicted. (**g**) Representative images of tumor burden, (**h**) spider plots, and (**i**) quantification of SB28 tumor burden following treatment. Two-way ANOVA with multiple comparisons, **p < 0.01, ***p < 0.001, ns = nonsignificant, mean ± s.e.m. depicted.

In a syngeneic SB28 glioblastoma model characterized by high MDSC infiltration (*48*), we evaluated whether sonothermogenetic control could enhance CAR T cell function by depleting MDSCs to promote a more favorable TME (**Fig. 6d**). Flow analysis revealed that the frequency of αEGFRvIII CAR T cells in tumors following sonothermal treatment were not significantly altered (**Fig. 6e**). However, sonothermal heating significantly reduced both the percentage and absolute count of tumor-infiltrating monocytic MDSCs (M-MDSCs) in the TS.mTCE +FUS cohort compared to unheated or WT controls (**Fig. 6f**). In a longitudinal study to assess therapeutic efficacy, WT T cells had no significant effect on tumor burden regardless of whether FUS treatment was used in combination, confirming that hyperthermia alone did not alter tumor progression. While cytotoxic TS.mTCE αEGFRvIII CAR T cells (**Fig. S8**) led to an initial reduction in tumor burden, they failed to sustain long-term tumor control. By contrast, FUS treated mice receiving TS.mTCE CAR T cell cohorts exhibited decreased tumor growth, with luminescence imaging showing markedly lower signals compared to all other control groups (**Fig. 6g-i**). These findings provide support that sonothermal activation of TS.mTCE αEGFRvIII CAR T cells can overcome intratumoral immunosuppression by *in situ* mTCE production to reduce the population of M-MDSCs.

## Discussion

Our study establishes closed-loop sonothermogenetic control to spatially and temporally regulate CAR T cell therapeutic activity within the brain. By integrating MRgFUS with CAR T cells engineered with thermal-sensitive bioswitches, we enabled metronomic production of NKG2D TCEs to overcome key barriers in brain tumor immunotherapy, including antigen heterogeneity and immunosuppression. Our findings establish that closed-loop MRgFUS can safely induce intracranial hyperthermia through the intact skull without lasting adverse effects, highlighting the translational potential of this platform. We validated this approach in multiple preclinical models, showing that localized TCE production by intratumoral CAR T cells redirected cytotoxicity towards antigen-negative tumor cells in a heterogeneous breast cancer brain metastasis model, and depleted MDSCs to remodel the glioblastoma microenvironment.

The selection of CAR T cell targets must balance specificity, efficacy, safety, and stability, which is an increasingly difficult task when targeting heterogeneous and immunosuppressive solid tumors such as glioblastoma with a single antigen(*49, 50*). To overcome this challenge our approach retains a tumor-specific CAR while introducing a second, switchable axis of cytotoxicity via NKG2D-based T cell engagers TCEs. Although this design enables both direct tumor cell killing and redirecting of bystander T cells to eliminate antigen-negative or stress ligand–expressing targets, the broad reactivity of NKG2D ligands increases the risk of off-tumor toxicity and cytokine release syndrome (CRS). By coupling TCE expression to a closed-loop sonothermogenetic control, we preserved the therapeutic advantages of dual-target CAR designs while integrating a layer of safety that is critical for clinical applications, where toxicity of “dual targeting” CAR T cells remains a major challenge (*2, 3*).

In the proposed research we employed clinical grade vectors derived from the humanized mAb Trastuzumab to engineer αHER2 CAR T cells. T cell engagers targeting CD19 are FDA-approved (Blinatumomab) for relapsed or refractory B-cell precursor acute lymphoblastic leukemia, and NKG2D ligands are targets in several ongoing clinical trials (NCT04658004, NCT04623944). Moreover, the application of trans-skull MRgFUS in the brain is approved by regulatory agencies for thermo-ablative interventions and our investigations are compatible with these systems (e.g., *ExAblate Neuro* by Insightec) (*32, 33*). It is also important to note that implantable US devices (e.g., SonoCloud-9 System by Carthera) could also be employed to test feasibility in GBM patients(*51*). The proposed closed-loop control methods (**Figs. 2–3**) can be readily integrated into these devices for application in humans for sonothermal activation of bioswitches with minimal adverse effects. Our approach can also be extended to enhance CAR T cell trafficking following intravenous delivery, using an additional sonication with the same device(*52*), to support a fully non-invasive and transcranial targeting of primary and metastatic brain tumors.

Collectively our research demonstrated the potential of closed-loop sonothermognetics to enable spatiotemporal control of CAR T cell therapeutic function in brain tumors. This approach offers new possibilities for metronomic immunotherapy and regulating the location, duration, and intensity of CAR T therapy to better manage toxicity and alleviate challenges associated with CAR T cell dysfunction, such as those mediated by persistent stimulation or an immunosuppressive TME. Together, our findings demonstrate that MRgFUS enables spatiotemporal control of CAR T cells, supporting the advancement of clinical translation of next-generation, remotely controlled cell therapies for treating brain tumors.

## Materials and Methods

### Ethics and Oversight

All research performed complied with relevant ethical regulations. Animal protocols were approved by the Georgia Tech Institutional Animal Care and Use Committee (IACUC). The use of recombinant DNA molecules or synthetic nucleic acid molecules were approved by the Georgia Tech Institutional Biosafety Committee (IBC).

### Statistics and reproducibility

Power analyses were performed using G*Power 3.1.9.6 (HHU) based on effect sizes determined by previous studies. Differences in animal survival (Kaplan-Meier survival curves) were analyzed by two-sided Mantel-Cox test. For statistical analysis of two groups, an unpaired Student’s t-test was applied. For grouped analyses, we performed a two-way ANOVA. Statistical analysis was performed using GraphPad Prism (Version 10.3.1). Data are presented as mean and s.d. or s.e.m. (error bars) as defined in the legends. Differences were considered significant when the p values were <0.05. Data distribution was assumed to be normal but this was not formally tested. Mice were inoculated with tumors and then randomized prior to allocation to experimental groups. For the remaining studies, randomization was not relevant, as these experiments were conducted on uniform biological materials, such as commercially sourced cell lines. The investigators were not blinded to group allocation during experiments and outcome analysis. Blinding was not feasible for both the in vitro and in vivo studies, as these experiments were conducted by individual investigators who were aware of the experimental groups. Additionally, blinding was not considered relevant, as the data analysis was performed quantitatively (e.g., flow cytometry, ELISA) and did not involve subjective, qualitative assessments. Mice were randomly assigned to experimental groups, but the experiments were not randomized, and the investigators were not blinded to group allocation or outcome assessment. No data were excluded from the analyses.

### Primary human T cell production

Peripheral blood was drawn from healthy human donors, as approved by the Georgia Tech and Emory University Institutional Review Boards (IRB #H20288). Peripheral blood mononuclear cells (PBMCs) were isolated using Lymphoprep density gradient medium (STEMCELL Technologies, 07801) and SepMate-15mL tube (STEMCELL Technologies, 85415), according to manufacturer’s instructions. CD3+ cells were then isolated using the EasySep Human CD3 Positive Selection Kit II (STEMCELL Technologies, 17851) and activated using Dynabeads (ThermoFisher, 11131D) at a 3:1 bead-to-cell ratio. Activated cells were cultured in complete human T cell media (hTCM; X-vivo 10 [Lonza #04-380Q], 5% Human AB serum [Valley Biomedical, HP1022], 10 mM N-acetyl L-Cysteine [Sigma A9165], 55 uM 2-mercaptoethanol [Sigma, M3148-100ML] supplemented with 50 U/mL recombinant human IL-2 (TECINTM Teceleukin, Bulk Ro 23-6019, National Cancer Institute, Frederick, MD) for 24 hours at 37°C in 5% CO2. To transduce the activated human T cells, concentrated lentivirus (MOI=30) was added to a 24-well suspension culture plate coated with Retronectin (Takara, T100B) according to the manufacturer’s instructions and subsequently centrifuged at 1,200×g for 90 minutes at 37°C. Following centrifugation, activated human T cells in human T cell media supplemented with 100 units/mL of hIL-2 were added to each well and the plate was spun at 1,200×g for 60 minutes at 37°C. Cells were incubated on the virus-coated plate for 24 hours before expansion. 7 days after activation, Dynabeads were removed.

### Primary murine CAR T cell production

Cells from C57BL/6 mouse spleens were harvested by gently dissociating the tissues using frosted glass slides, then centrifuged at 1000xg for 5 min and resuspended in 1x RBC lysis buffer for 5 min at 4°C. 1x PBS was added to quench the lysis reaction, and cells were centrifuged and resuspended in murine T cell media (mTCM; RPMI + 10% FBS + 1% Pen/Strep + 1x NEAA + 1x Sodium Pyruvate + 50 μM Beta-mercaptoethanol + 100 IU/mL rhIL-2) and passed through a 40 μm cell strainer. CD3+ cells from C57BL/6 mice isolated using the StemCell EasySep Mouse T Cell Isolation kit following the manufacturer’s protocol. After isolation, C57BL/6 CD3+ T cells were resuspended at a concentration of 1×10^6^ cells/mL in mTCM supplemented with murine Dynabeads at a 1:1 Bead to T cell ratio. Two days after activation, cells were collected, washed, and resuspended at 1×10^6^ cells/mL in concentrated retroviral supernatant supplemented with 100 IU/mL hIL2 and 8 μg/mL polybrene. Spinfection was performed in a U-bottom 96-well plate at 2,000xg for 90 min at 32°C. Transduced cells were maintained at 1×10^6^ cells/mL in complete murine T cell media supplemented with 100 IU/mL hIL-2 and passaged daily until use on Day 5.

### Brain tumor inoculations

SB28 glioma or MDA-MB-468 cells (0.5 - 2 × 10^4^ cells) were stereotactically implanted into the brains of 6-to 10-week-old female albino C57BL/6 J or NSG mice respectively (The Jackson Laboratory). The implantation site was located approximately 1 mm anterior and 1 mm to the right of the bregma. A small hole was drilled at the implantation site, and a syringe was inserted to a depth of 2.5 mm to inject the cells. The cells were injected via slow infusion over 5 minutes. Tumor growth was monitored using T2-weighted MRI and treatment occurred when bioluminescent signal reached sufficient signal. To minimize differences related to tumor size, before each experiment, the tumors in all animals were measured with MRI and IVIS and distributed equally among cohorts. The animals were considered as their endpoint if they exhibited severely impaired activity, significant weight loss, tumor dimensions exceeding 20 mm, or if treatment-related severe adverse events occurred that caused pain or distress and that could not be ameliorated.

### Adoptive cell transfer of CAR T cells

Transduced human CAR T cells were purified using the EasySep Human CD19 Positive Selection Kit II (STEMCELL Technologies, 17854) according to the manufacturer’s protocol after 9 days of activation. All Human T cells were maintained at a concentration of 0.7–2×10^6^ cells/mL until Day 10-14 for use in downstream assays. In contrast, murine T cell transduction was evaluated by GFP reporter expression 5 days post isolation. For ACT, 5 x 10^5^ CAR T cells were resuspended in 3 μL of sterile PBS and were transferred by stereotactic injection into brain tumors 1 mm anterior, 1 mm to the right, and 3 mm deep of the bregma of mice upon tumor engraftment assessed by IVIS or MRI.

### *In vivo* MRgFUS hyperthermia

Using frequency and design parameters optimized through mathematical modeling, we developed a custom-built closed-loop MRgFUS system for *in silico* validation, which we demonstrated in prior work(*30*). Briefly, the MRgFUS system operates at an optimized frequency of 1.7 MHz and comprises three key components: MR Thermometry, Transcranial FUS, and a Feedback Controller. The integration of transcranial FUS with MR thermometry allows precise monitoring and control of temperature elevation within the focal region. Additionally, the system incorporates a robust feedback control mechanism that leverages real-time MR thermometry to regulate thermal exposure, ensuring fine-tuned and sustained temperature control. Further details about the system and its validation are available in our prior study(*30*).

During hyperthermia treatments, we evaluated closed-loop feedback controllers (bi-state or PID) to implement various mild hyperthermia protocols (e.g., 10 minutes of hyperthermia at 41.5°C or 42.5°C using continuous wave (CW) or pulsed wave (PW) protocol). We rigorously tested the controllers through robust *in vivo* experiments, with a particular focus on the PID controller, which we introduced in this study. We optimized the PID controller’s proportional (P), integral (I), and derivative (D) values using the Ziegler-Nichols method and, in some cases, further refined these parameters through trial- and-error tuning to achieve effective thermal control. Temperature data extracted from a defined region of interest in the MR thermometry images informed the controllers, ensuring precise and reliable regulation of thermal exposure.

### Quantification of tumor-infiltrating immune cells and flow cytometry

To assess tumor composition, we performed flow cytometry on single cell suspension of tumor milieu. mice were euthanized and then immediately received intracardiac perfusion of 10 mL sterile PBS. The whole brain was dissected, and the entire tumor mass was separated from the brain. Tumors were immediately minced with razor blades and washed with RPMI. Tumors were dissociated using the Mouse or Human Tumor Dissociation Kits and gentleMACS Dissociator. Homogenized cells were passed through a 40 μm cell strainer and depleted of red blood cells using 1x RBC lysis buffer. TILs were isolated from the single cell suspension using a density gradient with Percoll Centrifugation Media and DMEM Media (10% FBS, 1% Pen-strep) at a 44:56 volume ratio. Cell viability was assessed by staining with LIVE/DEAD™ Fixable Aqua Dead Cell Stain Kit) following manufacturer’s instructions.

### Brain tissue processing

Following the treatments/sonications, the animals were euthanized and transcardially perfused with 20 ml of saline followed by 4% paraformaldehyde (PFA) before harvesting the brains. The brains were fixed with 4% PFA overnight at 4°C, followed by immersion in a 30% sucrose solution at 4°C until they sank to the bottom of the container. The brains were then embedded in O.C.T. compound and rapidly frozen to −80°C. Subsequently, 20 μm sections were cut using a cryostat (Leica 3050 S Cryostat) for further analysis.

### Immunofluorescence/Immunohistochemistry staining and microscopy

To assess the safety and the biological effects induced by FUS thermal stress, immunofluorescence staining was performed on brain tissue slides. Tissues were first fixed in 4% paraformaldehyde at room temperature for 10 minutes. For sections requiring staining of intracellular markers (e.g., GFAP and Iba-1, which are markers for astrocyte and microglial activation for safety assessment), they were subsequently permeabilized with 0.1% Triton X-100 in PBS for 5 minutes. After washing with PBS, the sections were blocked for 1 hour at room temperature in a solution containing 1% Bovine Serum Albumin and 5% goat serum in PBS.

The sections were then incubated with the primary antibody of interest, diluted in 1% Bovine Serum Albumin (1:100), for 12 hours at 4°C. Following primary antibody incubation, the sections were incubated with the secondary antibody, diluted in 1% Bovine Serum Albumin (1:200), for 1 hour at room temperature. To stain the cell nucleus, samples were incubated with DAPI diluted in PBS (1:1000) for 10 minutes after washing. Finally, the sections were rinsed with PBS to remove excess antibody, mounted with mounting medium, and covered with coverslips. Samples were cured with mounting medium for 24 hours in the dark at room temperature before imaging. The sections were imaged with a 20x objective using a fluorescence microscope (Eclipse Ti2, Nikon). The quantification of the fluorescence images was performed using ImageJ.

Hematoxylin-eosin staining was performed to examine tissue damage and safety. Twenty-micrometer thick frozen sections (Leica 3050 S Cryostat) were dehydrated beforehand and stained using a Leica Autostainer (ST5010). The sections were imaged with a 20x objective using a brightfield microscope (Eclipse Ti2, Nikon).

## Acknowledgements

We thank Dr. Hideho Okada (UCSF) for his helpful insights and contribution of SB28 and SB28-EGFRvIII cell lines.

## Author Contributions

Conceptualization: G.F.W. C.A., G.A.K.; Methodology: A.Z., C.K., C.A., G.A.K; Investigation: A.Z., C.K., S.S., S.F., R.S., N.C., S.A.O., Z.Z., C.T., E.K., M.B.; Visualization: A.Z., C.K.; Writing A.Z., C.K. C.A., G.A.K; Supervision: C.A., G.A.K.; Funding acquisition: C.A., G.A.K.; Project administration: G.A.K., C.A.

## Data and Materials Availability

All data needed to evaluate the conclusions in the paper are present in the paper and/or the supplementary materials. Material will be made available upon reasonable request to the corresponding author.

