## Supplement for "Closed-loop sonothermogenetic control of CAR T cells for metronomic brain cancer therapy"

**Fig. S1.**

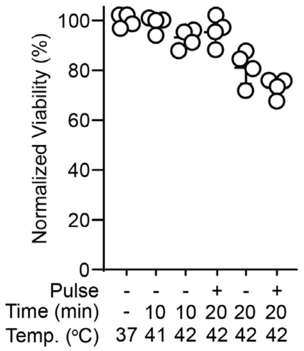


**Impact of FUS-mediated hyperthermia on primary human T cell viability.** Viability of TS.Fluc aHER2 CAR T cells following FUS mediated hyperthermia at indicated parameters quantified by flow cytometry.

**Fig. S2.**

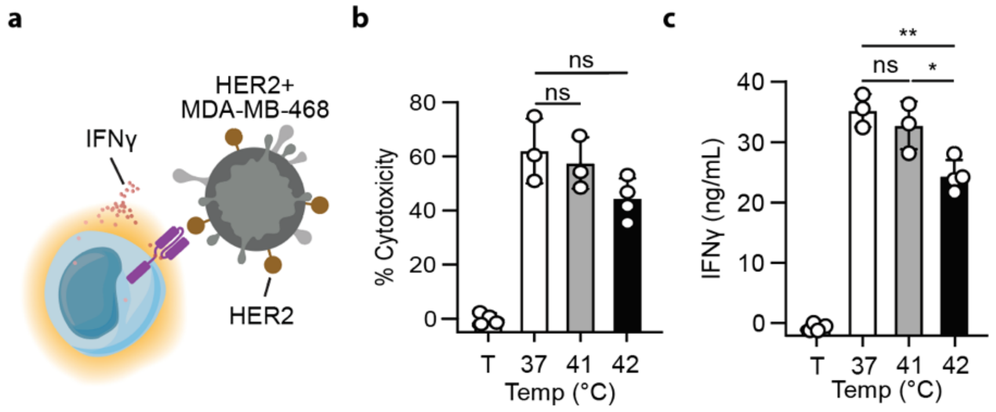


**FUS mediated hyperthermia at 41°C does not compromise key T cell functions.** (**a**) TS.Fluc aHER2 CAR T cells were heated at indicated temperatures for 20 minutes by FUS and then cocultured on HER2+ MDA-MB-468. (**b**) cytotoxicity and (**c**) IFNγ production was quantified 24 hours following coculture. One-way ANOVA with multiple comparisons, $*p < 0.05$, $**p < 0.01$, ns = nonsignificant, mean ± s.e.m. depicted, n= 3-4 independent wells, T= tumor only condition.

**Fig. S3.**

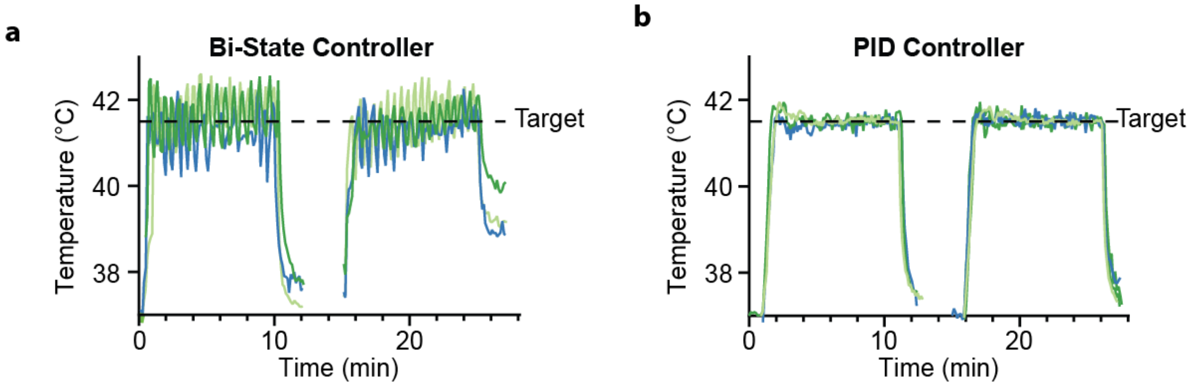


**Profile of temperature deposition varies by controller type.** Representative temperature profiles of 3 mice from (**a**) Bi-state controller and (**b**) PID controller closed loop system.

**Fig. S4.**

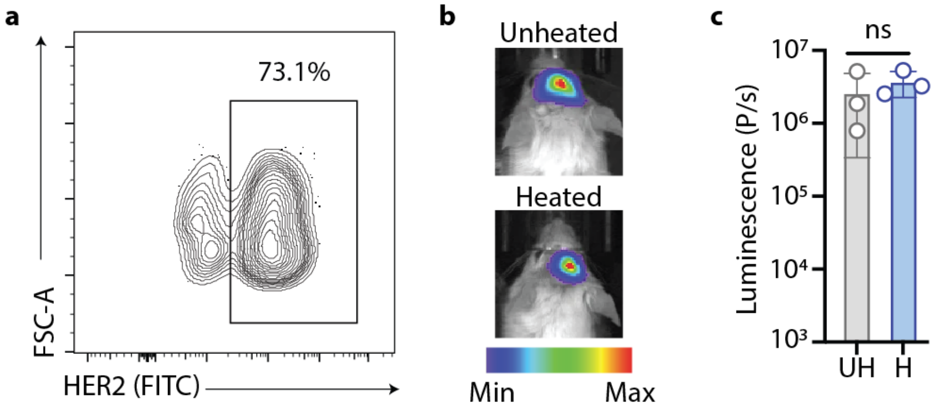


**FUS treatments do not alter tumor composition.** (**a**) A 3 to 1 mixture of HER2+/Fluc- to HER2-/Fluc+ MDA-MB-468 cells were inoculated into brains of NSG mice. (**b**) Representative IVIS image and (**c**) quantification of HER2-/Fluc+ tumors *in vivo* quantified 24 hours following thermal treatment. One-way ANOVA with multiple comparisons, ns = nonsignificant, mean ± s.d. depicted, n= 3 independent mice.

**Fig. S5.**

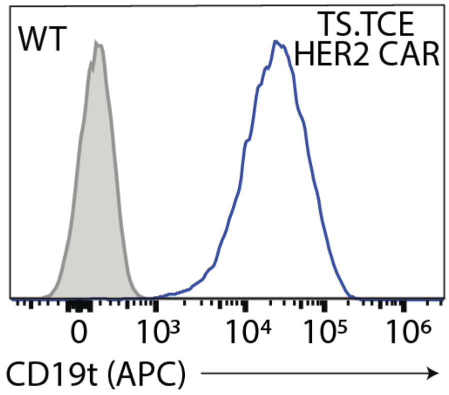


**Purification of transduced TS.TCE HER2 CAR T cells.** Flow cytometric validation of purity following MACS purification of effector TS.TCE HER2 CAR T cells.

**Fig. S6.**

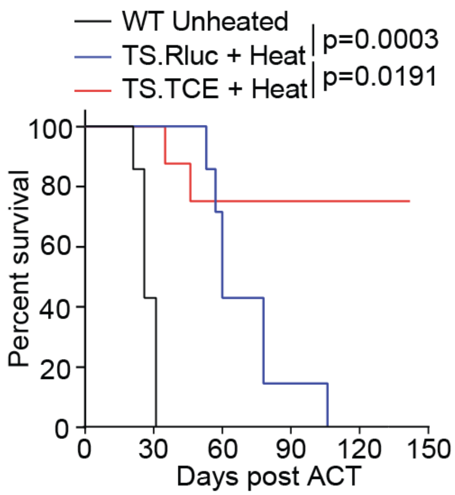


**Survival curves of heterogenous BCBM bearing mice**. Survival of tumor-bearing mice following treatment quantified for 150 days. log-rank (Mantel–Cox) test with P-values depicted.

**Fig. S7.**

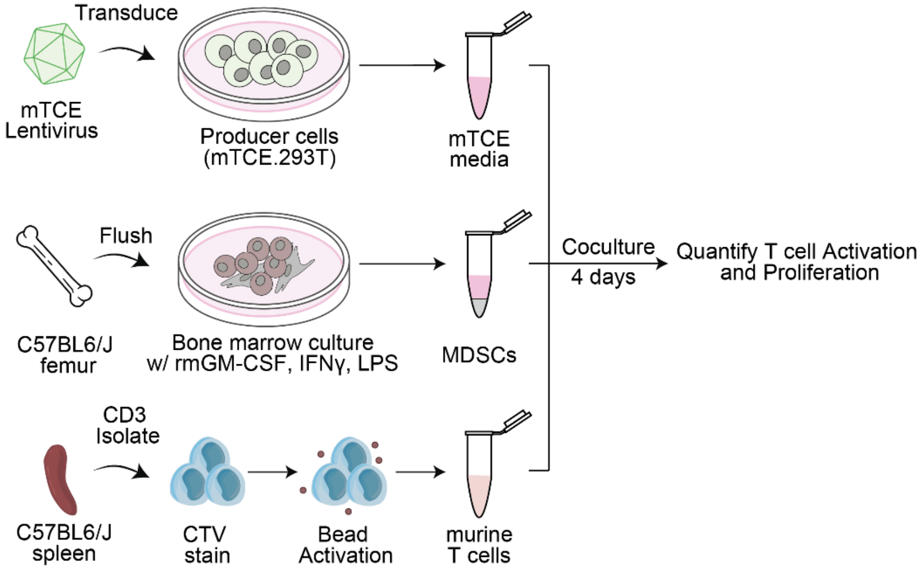


**Schematic representation of MDSC coculture assay.** mTCE producer 293Ts were generated by lentiviral transduction and cultured to generate mTCE conditioned media. Bone marrow derived MDSCs were differentiated using recombinant murine GM-CSF, IFNγ, and LPS. Splenic T cells were isolated, CTV stained and then activated by murine Dynabeads prior to coculture.

**Fig. S8.**

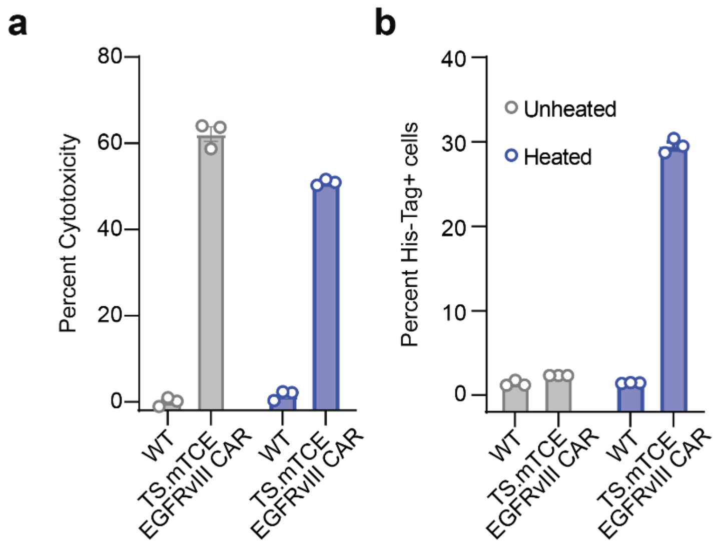


***In vitro* validation of TS.mTCE EGFRvIII CAR T cells. (a)** Cytotoxicity of WT or TS.mTCE EGFRvIII CAR T cells was evaluated against EGFRvIII+ SB28 tumor cells following 10 minutes of thermal cycler heating at 42°C. (**b**) TCE expression was quantified following heating by His-Tag staining on the surface of T WT or TS.mTCE EGFRvIII CAR T cells.
